# Structural Characterization of the PawL-Derived Peptide Family, an Ancient Subfamily of Orbitides

**DOI:** 10.1101/2021.07.18.452857

**Authors:** Colton D. Payne, Mark F. Fisher, Joshua S. Mylne, K. Johan Rosengren

## Abstract

Plants are an excellent source of bioactive peptides, often with disulfide bonds and/or a cyclic backbone. While focus has predominantly been directed at disulfide-rich peptides, a large family of small, cyclic but non-disulfide bonded plant peptides, known as orbitides, has been relatively ignored. A recently discovered subfamily of orbitides is the PawL-Derived Peptides (PLPs), produced during the maturation of precursors for seed storage albumins. Although their evolutionary origins have been dated, in-depth exploration of the family’s structural characteristics and potential bioactivities remains to be conducted. Here we present an extensive and systematic characterization of the PLP family. Nine PLPs were chosen and prepared by solid phase peptide synthesis. Their structural features were studied using solution NMR spectroscopy and seven were found to possess regions of backbone order. Ordered regions consist of β-turns, with some PLPs adopting two well-defined β-turns within sequences as short as seven residues, which are largely the result of side chain interactions. Our data highlight that the sequence diversity within this family results in equally diverse molecular scaffolds. None of these nine PLPs showed antibacterial or antifungal activity.

---

Backbone head-to-tail cyclic peptides fall into many different families and can be found across all domains of life.^1–3^ These peptides have been of particular interest for their structural stability and protease resistance conferred by the cyclization of their peptide backbones.^4, 5^ In addition, many cyclic peptides also contain one or more disulfide bonds, a feature that is critical to their three-dimensional structures and substantial stability.^6^

Historically, cyclic peptides were discovered and isolated from plants based on bioactivity.^7, 8^ As their biosynthetic origins became known, transcriptomics has been favored for discovery as well as examining the genetic origins and processing mechanisms behind some of the cyclic peptide families.^9–11^ Interestingly, several different head-to-tail cyclic peptide families share a similar processing and cyclization mechanism. These include the kalata-type cyclotides,^12^ cyclic knottins^13^ as well as the more recently discovered PawS-Derived Peptides (PDPs)^9^ and PawL-Derived Peptides (PLPs),^14^ all of which are proteolytically processed from precursor proteins by enzymes known as asparaginyl endopeptidases.^9, 12, 14^

Numerous members of the cyclotide, cyclic knottin and PDP families have had their 3D structures solved, and the features of these families are well established. Cyclotides and cyclic knottins adopt the cyclic cystine knot motif, an arrangement of disulfide bonds in which a ring like shape is formed by two disulfides, which brace the backbone of the peptide, with the knot ‘tied’ by the third disulfide bond threading through this ring.^15^ Bioactivities reported for members of these families include uterotonic,^16^ insecticidal,^17^ and antimicrobial^18^ activity, as well as protease inhibition.^19^ While most cyclotides are produced from dedicated precursor proteins,^20^ some examples are encoded within precursor proteins for seed storage albumins.^12^ This hijacking of an existing precursor protein already destined to be processed by asparaginyl endopeptidases, is shared by the PDP family.^11^ Most PDPs are cyclic peptides that adopt a single β-sheet separated by loops and bridged by a disulfide bond.^21^ The prototypic member of the family, SFTI-1, has nanomolar potency as a trypsin inhibitor,^22^ but little is known about the bioactivities and physiological roles of other members.

The 53 PLPs known to date are closely related to the PDPs, but lack disulfide bonds. Head-to-tail cyclic plant peptides without disulfide bonds are collectively referred to as orbitides, thus PLPs represent an evolutionary related subfamily of orbitides.^14^ The orbitides have a range of biosynthetic origins, but are typically five to ten residue peptides with sequences mostly comprised of hydrophobic amino acids. X-ray crystallography has demonstrated that orbitides can adopt ordered turns, with the peptide backbone forming a stabilizing hydrogen bond network; however the total number of resolved structures remains low.^23^ Some orbitides reported over the course of the past two decades presented antimalarial,^24^ antifungal^25^ or cytotoxic activities.^26^ This research has yet to be extended in great detail to the PLPs. With both the knotted cyclic peptides and PDPs subjected to extensive research regarding function and structure, PLPs remain the last of the four structural families of head-to-tail cyclic peptides sharing this unique processing mechanism to be extensively structurally and functionally studied. Here we describe seven new solution NMR structures of PLPs and put their structural features into the context of observations of other orbitides, and we examine them for potential antimicrobial bioactivity.

## RESULTS AND DISCUSSION

### Chemical synthesis and purification of PLPs

A large number of PLP sequences have recently been discovered within albumin precursors of the Asteraceae family, and 53 of these have been confirmed *in planta* via LC-MS/MS.^14, 27–29^ This served to establish the existence of the PawL-derived Peptide subfamily of orbitides. To further expand the structural and functional understanding of the PLP family, nine PLPs of varied sequence lengths and compositions were here chosen for study, PLPs −6, −13, −16, −22, −29, −31, −38, −42 and −46. Genes encoding each of these PLPs are known. The peptides are seven to ten amino acid residues in length, including an array of hydrophobic residues at varying locations (Figure 1). All contain a proto C-terminal Asp, and most have a proto N-terminal Gly residue typical of peptides cyclized by AEPs.^30^ However, PLPs −6, −13 and −42 differ with the N-terminal residue being Phe for PLP-6 and Thr for PLPs −13 and −42.

**Figure 1:**
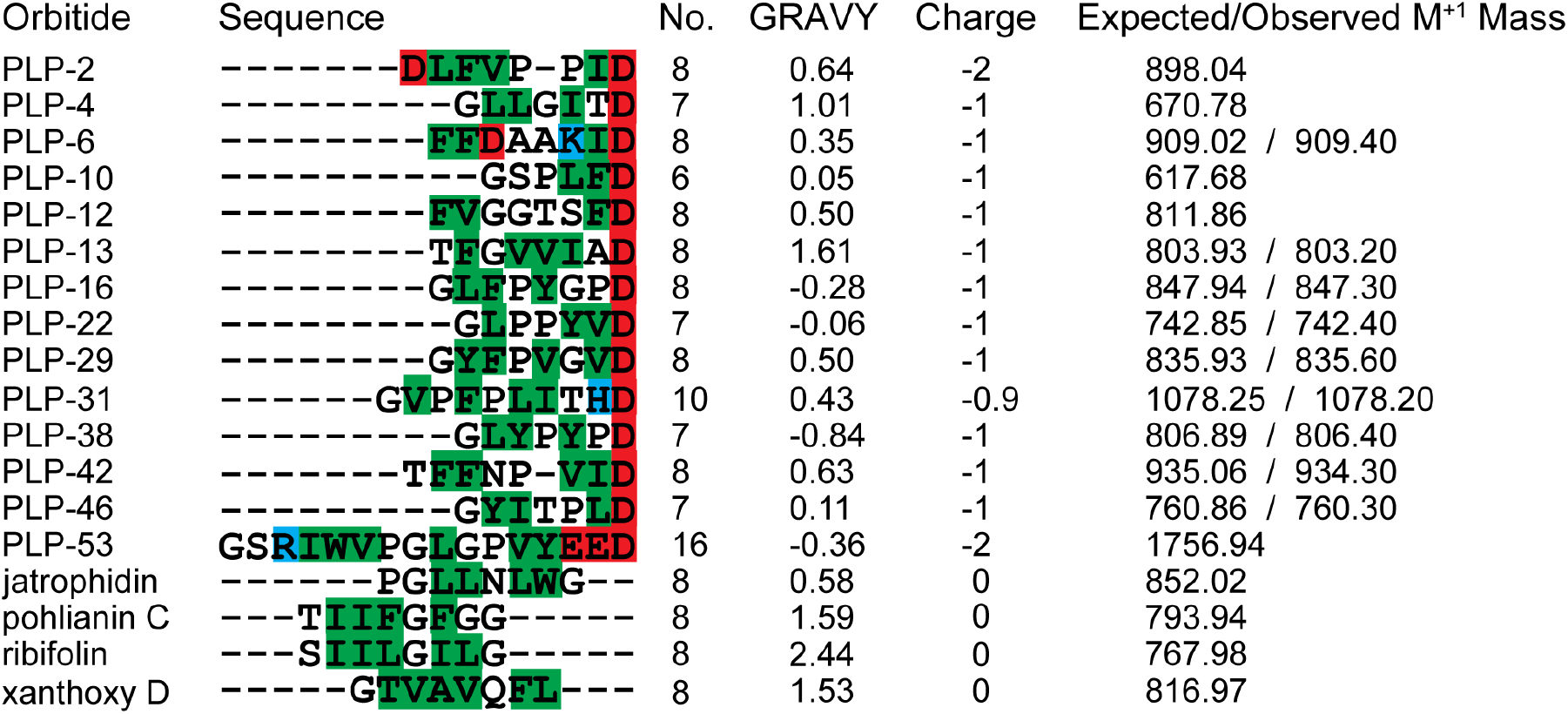
Alignment of PLPs and other orbitides, GRAVY scores, Net charge (pH 7) and expected/observed masses. PLPs chosen for this work (PLPs −6, −13, −16, −22, −29, −31, −38, −42 and −46) aligned with sequences for which structures have previously been solved (PLPs −2,^14^ −4,^14^ −10,^14^ −12^14^ and −53^29^ and the non-PLP orbitides jatrophidin,^23^ pohlianin C,^23^ ribifolin^23^ and xanthoxycyclin D^31^). Negatively charged residues (Asp and Glu) and positively charged residues (Arg, His and Lys) are highlighted in red and blue respectively. Hydrophobic residues are highlighted in green.

Peptides were assembled in full on resin using fluorenylmethyloxycarbonyl (Fmoc)-based solid phase peptide synthesis. After synthesis all PLPs were cleaved from the resin whilst maintaining side chain protection, and cyclized in solution before side chain deprotection. Likely due to the short sequence length of the PLPs, some peptides were prone to dimerization at high concentrations (10 mM) during the cyclization reaction. At low peptide concentration (1 mM) this was easily avoided, and effective cyclization achieved. Peptides were purified to >95 % purity by RP-HPLC and their identity confirmed by ESI-MS (Figure 1 in Supplementary Information).

### NMR spectroscopy of PLPs

To date, five PLPs have been studied by solution NMR spectroscopy, PLPs −2, −4, −10, −12 and −53.^10, 29^ PLP-2 was shown to have partially ordered structure, PLPs −4, −10 and −12 demonstrated a dynamic structure, and PLP-10 adopted at least two conformations based on proline isomerization. PLP-53, a uniquely large member of the PLP family with 16 residues in length, demonstrated a highly ordered region that had not been previously observed in the family. PLP-53 contains an extended turn region six residues in length, with the remaining ten residues forming a highly dynamic loop region.^29^ These findings are in contrast to the PDP family whose members, similar in size, are highly structured due to their disulfide bonds.^21, 32, 33^ To further understand the structural features adopted by the PLP family and potential key intramolecular interactions, PLPs −6, −13, −16, −22, −29, −31, −38, −42 and −46 were subjected to both homonuclear and heteronuclear solution NMR spectroscopy. As the majority of PLPs are comprised of almost entirely hydrophobic residues, with very few hydrophilic residues, many PLPs were insoluble in water. PLPs −6, −13, −22 and −31 were soluble in 90:10 H_2_O/D_2_O at different concentrations. However, PLPs −16, −29, −38, −42 and −46 were insoluble at any concentration in 90:10 H_2_O/D_2_O. NMR samples of these peptides were prepared in either 50:50 H_2_O/MeCN-*d*_3_ (PLPs −38 and −42), or 99.95% MeCN-*d*_3_ (PLPs −16, −29 and −46). The 1D spectra recorded for the nine PLPs varied in dispersion and sharpness, most peptides showed surprising dispersion of ^1^H NMR signals (Figure 2 in Supplementary Information). These features can be used to infer whether or not a peptide is adopting an ordered structure, with particularly wide dispersion of HN protons and sharp lines being strong indicators of a stable fold. Some PLPs have a markedly different 1D spectra to other members of the family. PLP-6 in particular presented HN signals with low dispersion and notably broad peaks, indicating disorder and either partial aggregation or intermediate conformational exchange. In addition, the ^1^H NMR spectrum of PLP-31 contained more signals than would be expected from its amino acid content, indicating the presence of multiple conformations in solution. For some of the PLPs few NOEs were observed, leading to issues resolving and sequentially assigning spin systems. Due to the low molecular weight of the PLPs this is not surprising as there is typically minimal NOE buildup in peptides of this size due to their fast tumbling in solution. As ROE buildup is more likely to be effective in these molecules ROESY data were also recorded for the PLPs. However, little to no difference was observed in the data between the NOESY and ROESY spectra. Reducing the temperature from 298K to 283K also had no significant effect on NOE buildup. However, sharp signals observed in the TOCSY and HSQC data allowed for the unambiguous assignment of almost all observed nuclei.

**Figure 2:**
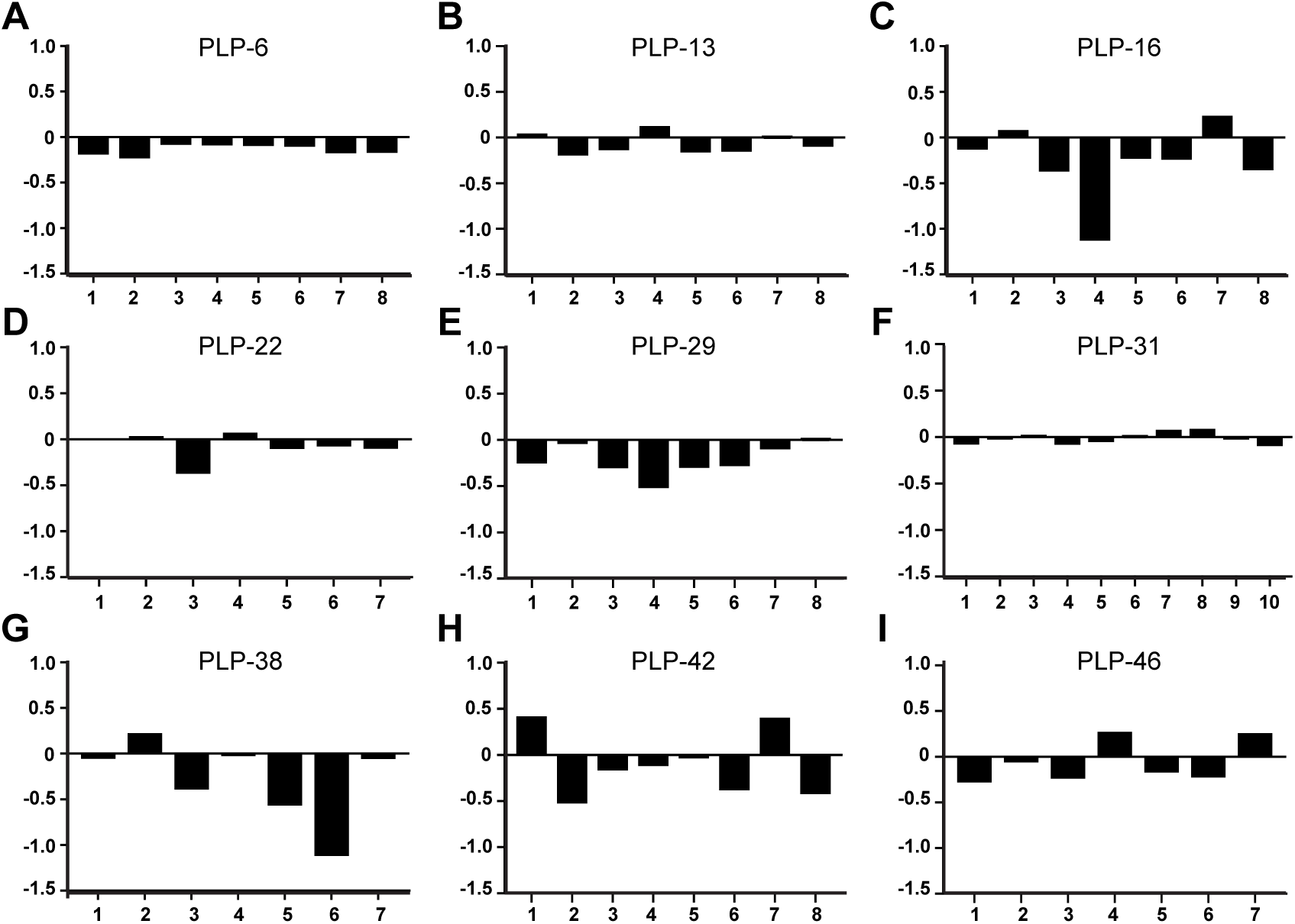
Secondary ^1^Hα chemical shifts of PLPs. A-I: PLP-6, −13, −16, −22, −29, −31, −38, −42 and −46. Strong deviations from random coil (large positive or negative values > 0.1 ppm) suggest order, whereas values close to zero suggest disorder. The y-axis for all graphs is the secondary chemical shift deviation (δΔ; ppm), the x-axis is the residue number.

The chemical shifts of Hα and Cα/Cβ nuclei are sensitive to backbone torsion angles and commonly these nuclei are analyzed to identify the presence of secondary structure in a peptide.^34, 35^ Thus, the Hα and Cα/Cβ chemical shifts for all PLPs were compared with random coil values (Figure 2 and Supplementary Figure 3). Chemical shifts that deviate substantially from the random coil values are indicative of ordered structure whereas values close to the random coil value are indicative of disorder. Typically, in larger peptides stretches of negative Hα secondary shifts are indicative of α-helical structure whereas stretches of positive deviation indicate β-sheet structure. Due to the short sequences of PLPs and the constrained cyclic backbone, it is unlikely that PLPs are able to adopt extensive secondary structure. Thus, these secondary shift graphs were used solely as a guide to indicate order or disorder. Hα secondary shifts with strong deviations from random coil were found in PLPs −16, −29, −38, −42 and −46. PLPs −6, −13 and −22 showed minor deviations, whereas PLP-31 appears essentially random coil throughout the sequence. Deviations from random coil were also found in the Cα/Cβ data for all PLPs, except PLP-31, which again appeared largely random coil. The HN, N, Hα, Cα and Cβ chemical shifts were used as input for the program Torsion Angle Likelihood Obtained from Shift and sequence similarity (TALOS-N)^36^ to predict the backbone dihedral angles of the PLPs. TALOS-N predicted a number of dihedral angles for seven of the PLPs (−13, −16, −22, −29, −38, −42 and −46) indicating ordered regions in those peptides. Notably, despite relatively small Hα secondary shifts, PLPs −13 and −22 were both predicted by TALOS-N to be ordered with a number of backbone dihedrals adopting defined angles. TALOS-N however makes predictions based on the combination of all backbone chemical shifts including those of sequential neighbors. This allows TALOS-N to confidently predict dihedral angles even if individual Hα/Cα/Cβ secondary shifts are not indicative of order. In contrast to PLP-13 and PLP-22, TALOS-N predicted that PLP-6 and PLP-31 are dynamic. This is consistent with the secondary shifts of PLP-31 and the poor dispersion of amide chemical shifts observed for PLP-6. As noted above, PLP-31 adopts several conformations in solution and the data used for this analysis only reflects the dominant and fully assignable conformation.

### Structure determination of PLPs

To evaluate the structural features of the PLPs, the 3D structures of PLP-13, −16, −22, −29, −38, −42, and −46 were determined using established protocols.^37^ PLP-6 and PLP-31 were not analyzed further given the multiple conformations of PLP-31 and the TALOS-N predictions of flexibility for both peptides. A combination of inter-proton distance restraints derived from NOESY cross peak volumes, backbone dihedral angle restraints determined by TALOS-N, hydrogen bond restraints based on temperature coefficients (Table 1 in Supplementary Information) and preliminary structure calculations were used. Preliminary structures were calculated using automated NOE assignment with CYANA 3.98.5.^38^ Final structures were calculated and refined in explicit solvent using CNS 1.21.^39^ All structures were analyzed using MolProbity^40^ to assess their stereochemical quality by comparison to published structures. The best 20 structures from the calculated 50 based on MolProbity scores, low energy and no significant violations of the experimental data, were chosen to represent the structural ensemble of each PLP (Figure 3, Table 1). All PLPs studied have good stereochemical quality with minimal to no Ramachandran outliers and atomic clashes. In addition, the backbone RSMD of all PLPs were below 0.3 Å in the ordered regions of the peptides. All PLPs have been submitted to the PDB and can be located under the following PDB and BMRB codes: PLP-13; 7M25 and 30881, PLP-16; 7M27 and 30882, PLP-22; 7M28 and 30883, PLP-29; 7M29 and 30884, PLP-38; 7M2A and 30885, PLP-42; 7M2B and 30886, PLP-46; 7M2C and 30887.

**Figure 3:**
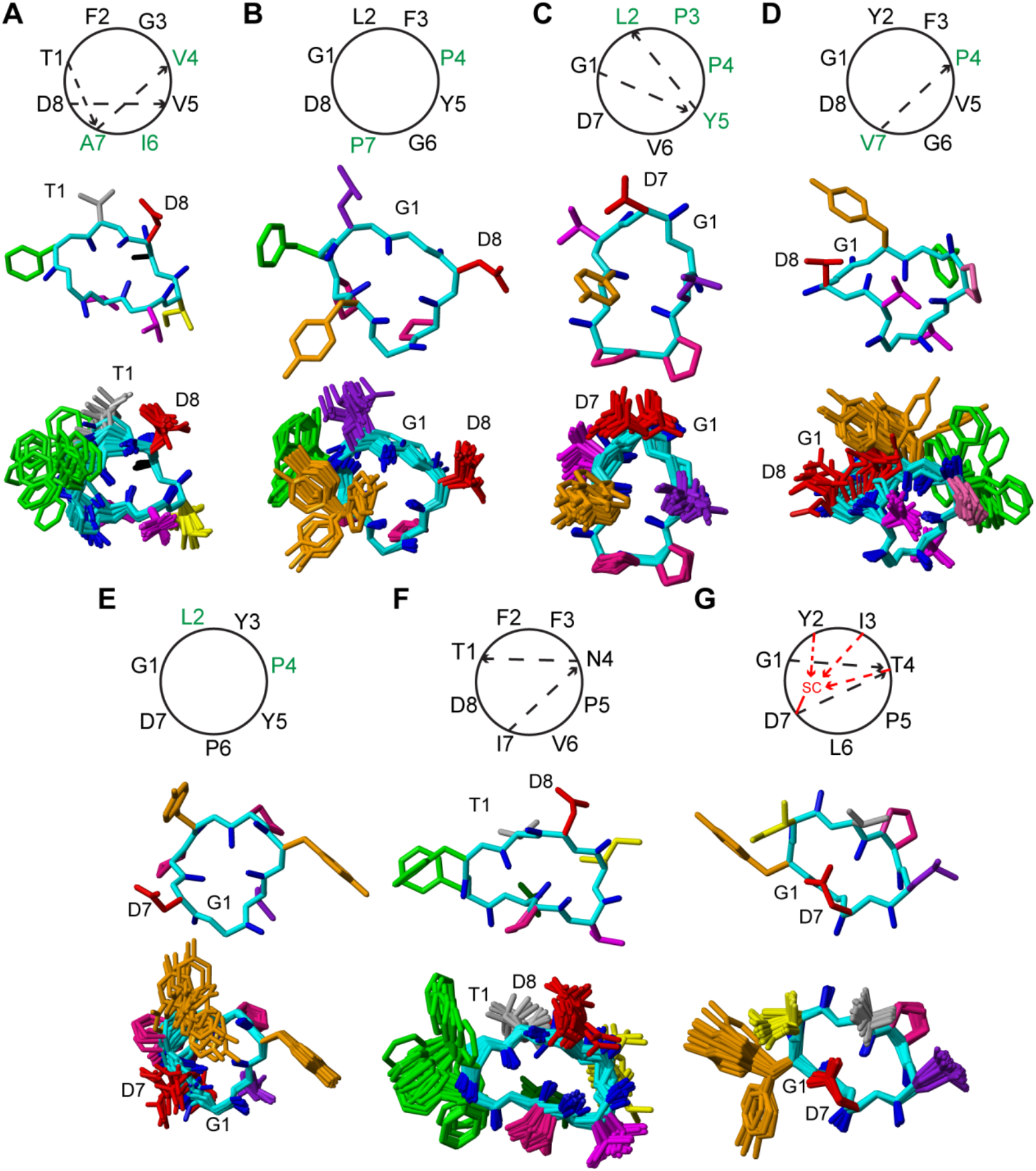
Hydrogen bond schematics and 3D structures of PLPs. A-G: PLP-13, −16, −22, −29, −38, −42 and −46. Hydrogen bonds are shown as dashed arrows (black arrows are from/to the peptide backbone, red arrows indicate hydrogen bonding to side chain groups). Green numbers indicate residues participating in hydrophobic side chain interactions. Three-dimensional structures are displayed with the backbone in cyan and the carbonyls in blue. Residues are colored as follows: Ala, black; Asn, dark green; Asp, red; Ile, yellow; Leu, purple; Phe, green; Pro, pink; Thr, grey; Tyr, orange; Val, magenta.

**Table 1:** Structure Calculation Statistics of PLPs.

| **Table 1: Structure Calculation Statistics of PLPs.** |  |  |  |  |  |  |  |
| --- | --- | --- | --- | --- | --- | --- | --- |
| **Energies (kcal/mol)** | **PLP-13** | **PLP-16** | **PLP-22** | **PLP-29** | **PLP-38** | **PLP-42** | **PLP-46** |
| Overall | -202.00<br>± 7.75 | -176.71<br>± 4.92 | -163.97<br>± 3.76 | -223.32<br>± 15.36 | -152.79<br>± 16.38 | -199.48 ±<br>14.73 | -245.72 ±<br>3.98 |
| Bonds | 2.25 ±<br>0.35 | 1.80 ±<br>0.23 | 2.13 ±<br>0.38 | 2.29 ±<br>0.39 | 2.59 ±<br>0.36 | 2.26 ±<br>0.39 | 2.30 ±<br>0.26 |
| Angles | 8.25 ±<br>1.62 | 5.34 ±<br>0.98 | 9.78 ±<br>1.28 | 9.60 ±<br>2.70 | 11.05 ±<br>2.64 | 10.81 ±<br>2.18 | 7.91 ±<br>1.04 |
| Improper | 3.16 ±<br>1.99 | 2.50 ±<br>1.26 | 2.06 ±<br>0.76 | 2.86 ±<br>1.23 | 5.03 ±<br>2.20 | 4.34 ±<br>1.87 | 2.74 ±<br>0.43 |
| Dihedral | 32.69 ±<br>0.56 | 25.32 ±<br>0.40 | 24.33 ±<br>0.41 | 28.05 ±<br>0.54 | 25.04 ±<br>0.52 | 34.98 ±<br>0.60 | 26.85 ±<br>0.42 |
| Van der Waals | -7.32 ±<br>3.15 | -17.89 ±<br>2.47 | -6.09 ±<br>2.08 | -17.46 ±<br>3.64 | -6.85 ±<br>2.76 | -14.11 ±<br>2.72 | -9.91 ±<br>2.26 |
| Electrostatic | -241.11<br>± 10.29 | -193.85<br>± 4.67 | -196.19<br>± 4.15 | -248.66<br>± 15.99 | -190.05<br>± 13.61 | -238.06 ±<br>15.88 | -275.99 ±<br>5.27 |
| NOE | 0.003 ±<br>0.004 | 0.002 ±<br>0.003 | 0.001 ±<br>0.002 | 0.003 ±<br>0.004 | 0.001 ±<br>0.008 | 0.001 ±<br>0.001 | 0.002 ±<br>0.002 |
| cDih | 0.067 ±<br>0.105 | 0.075 ±<br>0.089 | 0.003 ±<br>0.013 | 0.001 ±<br>0.005 | 0.381 ±<br>0.244 | 0.302 ±<br>0.285 | 0.395 ±<br>0.235 |
| **MolProbity Statistics** |  |  |  |  |  |  |  |
| Clashes (>0.4 Å/1000 atoms) | 0.88 ±<br>2.70 | 1.32 ±<br>3.21 | 0.00 ±<br>0.00 | 3.56 ±<br>5.32 | 10.65 ±<br>8.10 | 1.16 ±<br>2.84 | 0.00 ±<br>0.00 |
| Poor rotamers | 0 | 0 | 0 | 0 | 0 | 0 | 0 |
| Ramachandran outliers (%) | 0 | 0 | 3.00 ±<br>7.33 | 0 | 0 | 0 | 0 |
| Ramachandran favored (%) | 90.00 ±<br>8.38 | 89.17 ±<br>11.18 | 76.00 ±<br>12.31 | 86.67 ±<br>11.60 | 95.00 ±<br>8.89 | 100.00 ±<br>0.00 | 85.00 ±<br>8.89 |
| MolProbity score | 1.01 ±<br>0.35 | 1.04 ±<br>0.38 | 1.21 ±<br>0.26 | 1.33 ±<br>0.56 | 1.56 ±<br>0.58 | 0.64 ±<br>0.34 | 1.06 ±<br>0.33 |
| MolProbity score percentile | 99.00 ±<br>1.12 | 98.75 ±<br>1.33 | 98.6 ±<br>0.99 | 91.75 ±<br>13.01 | 86.30 ±<br>16.77 | 99.55 ±<br>1.10 | 99.25 ±<br>0.44 |
| **Atomic RMSD (Å)** |  |  |  |  |  |  |  |
| Mean global backbone | 0.66 ±<br>0.36 | 0.32 ±<br>0.11 | 0.40 ±<br>0.24 | 0.72 ±<br>0.39 | 0.66 ±<br>0.34 | 0.53 ±<br>0.24 | 0.16 ±<br>0.05 |
| Mean global heavy | 2.00 ±<br>0.73 | 1.75 ±<br>0.42 | 1.12 ±<br>0.30 | 1.86 ±<br>0.53 | 2.10 ±<br>0.71 | 1.96 ±<br>0.45 | 1.09 ±<br>0.47 |
| **Experimental Restraints** |  |  |  |  |  |  |  |
| NOE distances |  |  |  |  |  |  |  |
| Short range (i – j < 2) | 29 | 17 | 28 | 27 | 27 | 10 | 41 |
| Medium range (i – j < 5) | 10 | 1 | 7 | 2 | 1 | 2 | 3 |
| Hydrogen bond restraints | 6 (3 bonds) | 0 | 2 (1 bond) | 2 (1 bond) | 0 | 4 (2 bonds) | 10 (5 bonds) |
| Total | 45 | 18 | 37 | 31 | 28 | 16 | 54 |
| Dihedral Angle Restraints |  |  |  |  |  |  |  |
| φ | 3 | 2 | 1 | 1 | 3 | 5 | 3 |
| ψ | 3 | 4 | 2 | 2 | 3 | 6 | 4 |
| Total | 6 | 6 | 3 | 3 | 6 | 11 | 7 |
| **Violations** |  |  |  |  |  |  |  |
| NOE violations > 0.2 Å | 0 | 0 | 0 | 0 | 0 | 0 | 0 |
| Dihedral violations > 2.0° | 0 | 0 | 0 | 0 | 0 | 0 | 0 |

### Structural features of PLPs

PLPs, like other orbitides, are too small and constrained by their cyclic backbone to adopt secondary structures like α-helices and β-sheets. However, this does not mean that PLPs are unable to adopt any form of structural order. The small size of the PLPs enforces turns in which, hydrophobic interactions and intramolecular hydrogen bonds can stabilize the defined backbone conformations. These features were observed in all the PLPs for which 3D structures were calculated.

PLP-13, which was studied in water, is one of the more ordered members of the PLP family with two overlapping turns and three backbone hydrogen bonds. Two of the observed hydrogen bonds stabilize a type III β-turn comprising residue 4-7 and a type I β-turn comprising 5-8. These well-defined features established a global RMSD of 0.66 Å (Figure 3A). Defining angles of each type of β-turn are supplied in Supplementary Table 2 for clarity. The remaining hydrogen bond is between the HN proton of Thr1 and the backbone carbonyl of Ala7. For proteins studied in aqueous solution, the presence of hydrogen bonds can be experimentally supported by monitoring the temperature dependence of amide proton chemical shifts.^41^ These hydrogen bonds are all consistent with the amide temperature coefficients determined for PLP-13 (Table 1 in Supplementary Information). The side chains of residues Ile6 and Ala7 participate in hydrophobic interactions with Val4, likely assisting in the stabilization of the two turns present in this region. Residues Phe2 and Gly3 are less defined and predicted by TALOS-N to be flexible. The side chains of Thr1, Phe2 and Asp8 are fully solvent-exposed.

Interestingly, no backbone hydrogen bonds were predicted for PLP-16. An atypical turn is formed by residues 4-7, based on backbone angles and the distance between Cα atoms. This is likely facilitated by the proline residues at positions 4 and 7. These two prolines, one of which is in a *cis* conformation (Pro4) and the other in a *trans* conformation (Pro7), also form a hydrophobic interaction with each other across the turn to further stabilize the backbone. Overall, this establishes a global RMSD of 0.32 Å (Figure 3B). Despite being large and hydrophobic, the remaining side chains do not appear to participate in any interactions and are largely solvent-exposed. It should be noted that the data for PLP-16 were recorded in 99.95% MeCN-*d*_3_ due to its poor solubility in water.

Two type III β-turns are formed in PLP-22 in H_2_O, one comprising residues 2-5 and the other residues 5-1, each supported by a backbone hydrogen bond, one of which was confirmed by the amide temperature coefficients (Table 1 in Supplementary Information). These turn regions are well-defined and establish a global RMSD of 0.40 Å (Figure 3C). PLP-22 contains two sequential Pro residues at positions 3 and 4, both in the *trans* conformation. Interestingly, all amino acid side chains, barring that of Val6, are projected on the same face of the PLP forming a densely packed hydrophobic surface.

PLP-29 only adopts only a single turn. A hydrogen bond between residues Pro4 and Val7 and hydrophobic contacts between the side chains of these residues stabilize a type II turn in this region. The remainder of the backbone of PLP-29 is less defined, but the backbone folds back on top of itself, and achieves an RMSD of 0.72 Å (Figure 3D). The remaining side chains, which are largely hydrophobic, project towards the solvent. Aromatic rings, when adjacent to a proline residue, can favor formation of a *cis* proline. In this case, the NMR data indicate that Phe3 does not interact with Pro4 and that Pro4 is in the *trans* conformation.^42^ Again, PLP-29 was studied in 99.95% MeCN-*d*_3_ because of its poor solubility in H_2_O. It cannot be ruled out that more defined packing of these side chains would occur in aqueous solution.

PLP-38 presents an interesting case wherein no hydrogen bonds were predicted and no turn region was observable. However, hydrophobic interactions between the side chains of residues Leu2 and Pro4 likely contribute to local order. Like PLP-16, the Pro4 and Pro6 residues in PLP-38 adopt *trans* and *cis* conformations, respectively. Despite a lack of hydrogen bonding, the global backbone RMSD of 0.66 Å suggests that PLP-38 has a stable conformation (Figure 3E). The side chains are solvent exposed in 50:50 H_2_O/MeCN-*d*_3_.

PLP-42, like PLP-13, is a highly ordered member of the family with two hydrogen bonds stabilizing two consecutive turns: A type I turn is formed by residues 1-4 and a type III turn is formed by residues 4-7. This double turn creates highly ordered regions and establishes a global backbone RMSD of 0.53 Å, only allowing for some dihedral angle ambiguity at Thr1 (Figure 3F). No notable side chain interactions were present, with most side chains being solvent-exposed based on the data recorded in 50:50 H_2_O/MeCN-*d*_3_.

PLP-46 is the most robustly stabilized member of the PLP family studied to date. PLP-46 contains five hydrogen bonds that fully constrain the backbone of the peptide, achieving a global backbone RMSD of 0.16 Å (Figure 3G). Two of these hydrogen bonds between residues Gly1 and Thr4 and Thr4 and Asp7 form a double turn comprised of a type IV and a type III turn respectively, similar to what was observed for PLP-42. The three remaining hydrogen bonds are formed between the backbone HN of residues Tyr2, Ile3 and Thr4 and the side chain carboxyl oxygens Oδ1 and Oδ2 of Asp7. This places the side chain of Asp7 in a unique location, almost centralized perfectly over the cyclic ring of the backbone, allowing these protons to form hydrogen bonds with the side chain. All other side chains lack notable interactions and are solvent-exposed in 99.95% MeCN-*d*_3_.

### How structurally similar are PLPs to other orbitides?

CyBase,^43^ a database of cyclic protein sequences, currently includes a total of 197 orbitide sequence entries, including the recently established 53 PLP sequences. Of these, eight orbitides have had their 3D structures solved by solution NMR spectroscopy or X-ray crystallography and the coordinates deposited in the Protein Data Bank (PDB). Nine orbitides have had structures solved by X-ray crystallography and deposited in the Cambridge Structural Database.^23^ Of the published orbitide structures, all contain at least one hydrogen bond stabilizing the backbone. Most of these hydrogen bonds stabilize turns, although some orbitides, like the PLPs studied here, contain other backbone-to-backbone or backbone-to-sidechain hydrogen bonds that further stabilize the peptide.

Intriguingly, most previously reported PLPs studied by NMR spectroscopy do not appear to adopt ordered structure in solution. Exceptions to this are PLP-2 and PLP-53. PLP-2 contains some order in a turn region. However, this is confined to three residues and likely due to two sequential proline residues locking its backbone configuration. The unusually large PLP-53 contains an extended turn region six residues in length with supporting hydrogen bonds.^29^ Two peptides studied in this work, PLP-6 and PLP-31, were found to have dynamic structures based on chemical shifts, and thus are consistent with the structures of most previously reported PLPs. The remainder of the PLPs studied in this work, however, adopted more ordered structures consistent with what was observed throughout the rest of the orbitide family. PLPs, despite their small size, can adopt one or several highly ordered turns stabilized by hydrogen bonds and hydrophobic side chain interactions. These features are difficult to predict based on sequence alone. It is interesting to note that PLP-53, the largest reported PLP at 16 residues in length, only adopts a singular turn region, while PLPs half the size, such as PLPs −13, −22, −42 and −46, adopt two turns. This work highlights that PLPs commonly adopt one or two ordered turn regions stabilized by hydrogen bonds and/or hydrophobic interactions, akin to other orbitides, further highlighting the PLPs position as a subfamily of the orbitides (Figure 4).

**Figure 4:**
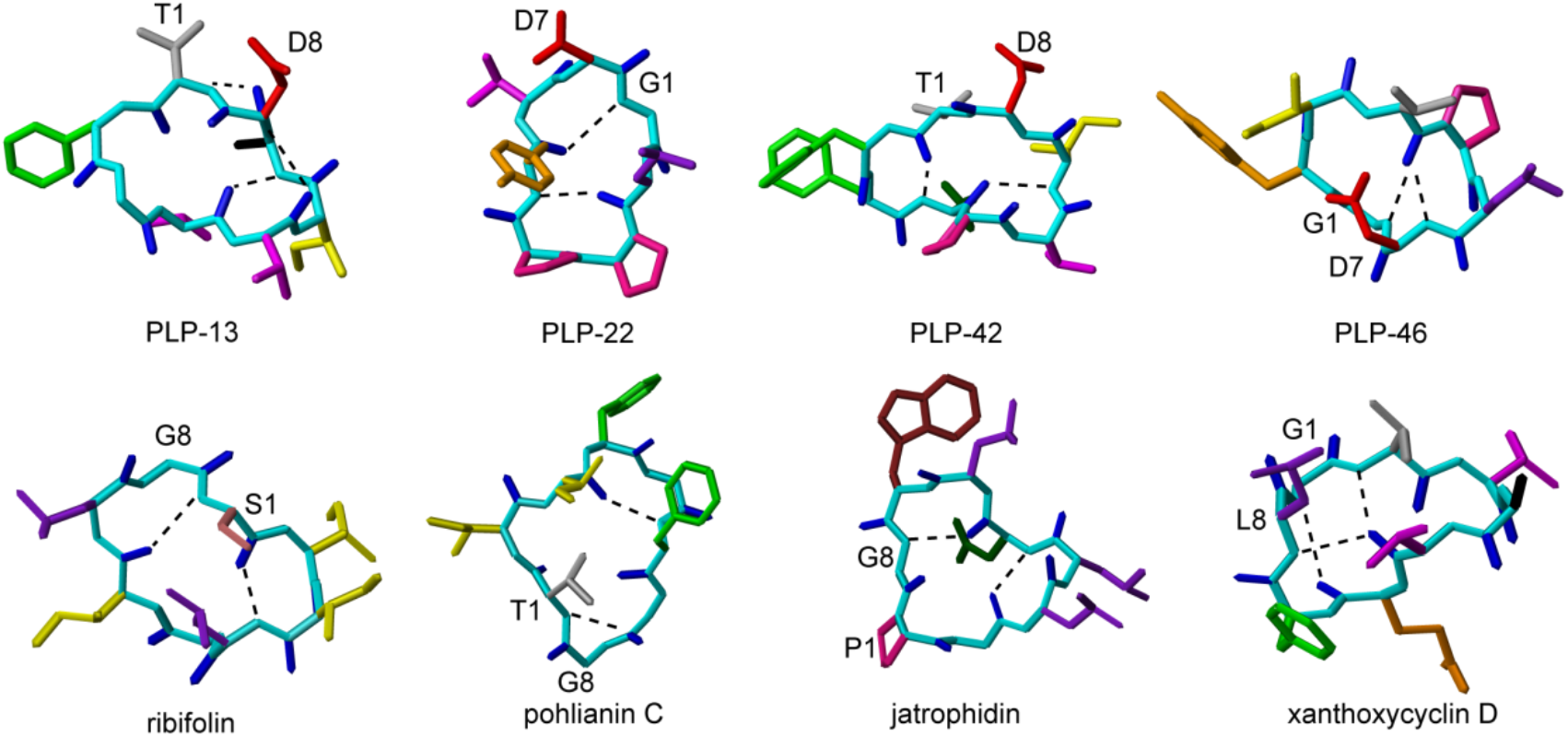
3D structures of PLPs compared to other orbitides. Peptides are labeled beneath structures and are colored as described in Figure 3 with the additions of Gln in gold and Trp in brown. Backbone hydrogen bonding is shown with dashed lines. PLPs −13, −22, −42 and −46 are from this work. Ribifolin (6DKZ) from *Jatropha ribifolia*,^23^ pohlianin C (6DL0) from *J. pohliana*,^23^ jatrophidin (6DL1) from *J. curcas*^23^ and xanthoxycyclin D (6WPV) from *Melicope xanthoxyloides*.^31^

### Are PLPs antimicrobial?

Since antifungal activity has been reported for orbitides, we tested our nine peptides for antifungal and antibacterial activity in a disc diffusion assay. Discs containing 200 μg of each

PLP were placed on YPD (yeast extract, peptone and dextrose) agar plates swabbed with *C. albicans* and LB agar plates swabbed with either *Escherichia coli* K-12 or *Bacillus subtilis* Marburg No. 165. None of the PLPs tested inhibited the growth of bacteria (Figure 4 in Supplementary Information). The effect on *C. albicans* was also minimal PLP-29 did show minor growth inhibition (Figure 5 in Supplementary Information). However, it was not possible to quantify the zone of inhibition since some fungal colonies were still present.

Given the potential activity of PLP-29 against *C. Albicans*, we performed a solution assay to determine its minimum inhibitory concentration. No measurable inhibition of growth was however observed at concentrations up to 640 μM (Table 3 in Supplementary Information). This result shows that PLP-29 is unlikely to have a useful antimicrobial function.

These results are consistent with a previous study which found that PLPs −2, −18 and −20 were not active against the Gram-positive bacteria *Bacillus cereus* and *Staphylococcus aureus*, nor against the Gram-negative bacteria *E. coli* and *Pseudomonas aeruginosa*.^14^ Looking at the structures and sequences of the tested orbitides, they have little in common with most antimicrobial peptides, which tend to be positively charged, allowing interactions with bacterial membranes.^44^ In contrast, all orbitides studied here carry a net negative charge. The orbitide tunicyclin D has potent antifungal activity against *C. albicans*.^25^ This peptide, lacks negatively charged residues but does contain a Trp and a His residue, which may allow membrane interactions leading to this bioactivity.

In summary, in this work we have synthesized and characterized nine PLPs. None were found to inhibit the growth of fungi or bacteria, leaving their physiological roles yet to be understood. Structural characterization by NMR spectroscopy resulted in the determination of seven new structures displaying highly ordered turn regions. The remaining two (PLP-6 and PLP-31) have a dynamic backbone. This work highlights that, like other orbitides, PLPs are restrained by backbone hydrogen bonds and side chain interactions. We show that the nature of these is diverse, highlighting the difficulty of predicting structure and dynamics of members of this family based on sequence alone.

## EXPERIMENTAL SECTION

### General Experimental Procedures

One-dimensional as well as 2D ^1^H-^1^H TOCSY^45^ and NOESY^46^ experiments with mixing times of 80 and 400 ms, respectively, were recorded at 298 K on a 900 MHz Bruker Avance III spectrometer with a cryoprobe. TOCSY experiments were recorded with 8 scans and 512 increments, NOESY experiments were recorded with 24-40 scans and 512 increments depending on signal-to-noise in the 1D spectra. Both TOCSY and NOESY data were recorded with a sweep width of 10 ppm due to the lack of Trp residues in PLP sequences. ^1^H-^13^C and ^1^H-^15^N HSQC spectra were recorded at 298 K at natural abundance. The ^1^H-^13^C experiments were recorded with 72 scans and 256 increments with a 10 ppm sweep width in the F2 dimension and an 80 ppm sweep width in the F1 dimension. The ^1^H-^15^N experiments were recorded with 128 scans and 128 increments with a sweep width of 10 ppm in the F2 dimension and 32 ppm in the F1 dimension. TOCSY temperature data were also recorded for all PLPs at 288, 293, 298, 303 and 308 K to monitor the temperature dependence of the amide protons. Data processing was conducted using Topspin 4.0.3 (Bruker) with the spectra being referenced to the solvent signal (spectra recorded at 298 K were referenced to the residual solvent at 4.769 ppm for H_2_O/D_2_O data and 2.031 for MeCN-*d*_3_ data). All PLPs were purified using reverse phase HPLC with a Prominence (Shimadzu, Rydalmere, AUS) instrument using a C18 preparative column (300 Å, 10 μm, 21.2 mm i.d. x 250 mm, Phenomenex). Analysis by ESI-MS was used to confirm the mass of the PLPs synthesized (Figure 1 in Supplementary Information). Further purification was conducted by HPLC using a semi-preparative C18 column (300 Å, 5 μm, 10 mm i.d. x 250 mm, Vydac). Final purity was assessed with a C18 analytical column (300Å, 5 μm, 2.1 mm i.d. x 150 mm, Vydac). All purifications were conducted using an increasing 1%/min gradient of 90:10 MeCN/H_2_O, with all PLPs confirmed to be >95% pure (Figure 1 in Supplementary Information).

### NMR Spectroscopy

NMR samples were prepared by dissolving 1-2 mg of peptide in 500 μL of solvent at pH ~3.5. PLPs −6, −13, −22 and −31 were dissolved in (90:10) H_2_O/D_2_O, PLP-38 and PLP-42 were dissolved in (50:50) H_2_O/MeCN-*d*_3_ and PLPs −16, −29 and −46 were dissolved in 99.5% MeCN-*d*_3_. Sequential assignment strategies^47^ were used to analyze all TOCSY and NOESY data; heteronuclear data were analyzed with the assistance of assigned TOCSY/NOESY data within the program CARA.^48^ Secondary Hα/Cα/Cβ shifts were calculated via shift comparison to random coil values to determine order, if any, within the PLPs.^34^

### Peptide synthesis

PLPs were synthesized using Fmoc based solid phase peptide synthesis on 2-chlorotrityl resin at a 0.125 mmol scale. Loading of the C-terminal residue was achieved by dissolving 1 M eq. of the Fmoc-protected residue in minimal CH_2_Cl_2_ and 4 M eq. of N,N’-diisopropylethylamine (DIPEA) before addition of this solution to 2-chlorotrityl resin pre-swollen in CH_2_Cl_2_ for 1 h. Standard deprotection was conducted between couplings using 2 x 5 min reactions of 20% v/v piperidine in DMF. Subsequent amino acids were added to the peptide chain by activating 4 M eq. of the Fmoc protected amino acid in 4 M eq. of 0.5 M 2-(1H-benzotriazole-1-yl)-1,1,3,3-tetramethyluronium hexafluorophosphate and 8 M eq. of DIPEA and applying the solution to the resin using a CS336X peptide synthesizer (CSBio, CA, USA). Couplings were conducted twice for all amino acids that contain branched β-carbons or aromatic rings. Upon completion of synthesis, peptides were cleaved from the resin using 10 x 3 min treatments of 2% (v/v) trifluoroacetic acid (TFA) in CH_2_Cl_2_ to maintain side chain protection. MeCN was added to the cleavage solution and rotary evaporation was conducted to remove the CH_2_Cl_2_ and TFA, with the remaining solution being lyophilized. Cyclization of the side chain protected PLPs was conducted in DMF at a 1 mM peptide concentration, 1 M eq. of 1- [bis(dimethylamino)methylene]-1H-1,2,3-triazolo[4,5-b]pyridinium 3-oxide hexafluorophosphate was added to this solution followed by the slow addition of 10 M eq. of DIPEA. The reaction was monitored over 2 h using electrospray ionization mass spectrometry (ESI-MS). Phase separation with CH_2_Cl_2_ was used to isolate the peptide from the DMF after the reaction was completed. The CH_2_Cl_2_ solution containing the peptide was diluted with MeCN, evaporated and the peptide solution was lyophilized. PLPs had side chain protection removed using a 97:2:1 TFA/H_2_O/triisopropylsilane solution. The solution was stirred for 2 h before precipitation of the peptide using cold Et2O. The precipitate was filtered and re-dissolved in 50:50 MeCN/H_2_O before lyophilization.

### Structure calculation

Inter-proton distance restraints were generated from the peak volumes of cross peaks present in the NOESY spectra for each PLP. TALOS-N^36^ was used to predict dihedral ϕ (C_-1_-N-Cα-C) and ψ (N-Cα-C-N_+1_) backbone angles. Potential hydrogen bond donors were identified from backbone amide temperature coefficients. Values > −4.6 ppb/K for the coefficient of the linear relationship were taken to indicate a hydrogen bond donation by the particular HN proton.^41^ Hydrogen bond acceptors were determined through preliminary structure calculations. Initial structures (50) were calculated in CYANA 3.98.5^38^ using torsion angle simulated annealing, which defined the starting coordinates and distance restraints to be used in the CNS.^39^ TALOS-N dihedral restraints, distance restraints from CYANA and hydrogen bonds from temperature coefficients were input into CNS 1.21.^39^ A further 50 structures were generated in CNS by simulated annealing using torsion angles.^39^ The annealed structures were then subjected to water minimization using Cartesian dynamics to generate final PLP structures. Stereochemical analysis was conducted by MolProbity via the comparison of the generated structures to previously published structures from RCSB.^40^ MOLMOL^49^ was used to display and generate images of structural ensembles with the 20 best structures, defined as having good geometry, no violations of distances of dihedral angles above 0.2 Å or 2° and low energy.

### Antimicrobial disc diffusion assays

The nine peptides were each dissolved in DMSO to a concentration of 20 mg/mL for disc diffusion assays carried out using an existing protocol,^14^ modified such that only one dose of each peptide was used. For the antibacterial assays, cultures of *E. coli* K-12 and *B. subtilis* Marburg No. 165 in liquid LB medium were diluted to an OD_600_ of 0.1 and spread onto LB agar plates with a sterile swab. For antifungal assays with *C. albicans*, the same procedure was used except that a YPD culture was diluted to an OD_600_ of 0.05 and spread onto YPD agar plates.

Once the cultures had been absorbed by the medium, we dispensed 10 μL (200 μg) of each peptide onto sterile 8 mm filter paper discs, allowed them to dry for 2 h in sterile conditions and placed them on the inoculated plates. We also used discs containing 5 μL DMSO as a negative control on each plate. Positive controls were discs containing 50 μg of kanamycin for the antibacterial assays or 50 μg of amphotericin B for the antifungal assays. We hypothesized that the presence of TFA counterions in the peptide samples (due to chemical synthesis) might have an effect on the assay. To control for this we removed any traces of TFA from all the peptides, via salt exchange with HCl, prior to dissolving them in DMSO. One aliquot of PLP-13 was left untreated for the purposes of comparison. The cultures were incubated at 37 °C for 16 h (bacteria) or 20 h (fungus), then inspected for any inhibition of microbial growth.

### Antifungal minimum inhibitory concentration assay

To confirm the antifungal activity of PLP-29, we performed an assay to determine its minimum inhibitory concentration was performed. Onto a 96-well plate 200 μL aliquots of *C. albicans* cultures containing PLP-29 were dispensed to a final concentration of 5, 10, 20, 40, 80, 160, 320 or 640 μM in three replicates. The aliquots were composed of 180 μL of fungal culture in YPD medium adjusted to an OD_600_ of 0.05 and 20 μL of peptide solution. The peptide solution was prepared by diluting an initial solution in DMSO at 20 mg/ml with H_2_O to a concentration of 6.4 mM (5.9 mg/ml), and then preparing serial dilutions in H_2_O for the lower concentrations. Further cultures containing amphotericin B as a positive control were dispensed in the same concentrations, again in three replicates. A series of negative control cultures contained DMSO to the same concentrations as the PLP-29 and amphotericin B samples (both amphotericin B and PLP-29 were initially dissolved in DMSO before further dilution with H_2_O). A “no inoculum” series of controls contained growth medium and PLP-29 only, in the same concentrations as above. After measuring the initial OD_595_ of each well with a FLUOstar OPTIMA plate reader (BMG Labtech GmbH, Offenburg, Germany), the plate was incubated for 7 h at 37 °C, after which the final OD_595_ was measured. A longer time course was also attempted, but over 24 h the fungi grew too much to reliably measure the optical density of the culture. Fungal growth was measured by the change of optical density over time, comparing the wells containing DMSO only with those containing either PLP-29 or amphotericin B.

## Author Information

## Author Contribution

C.D.P.: peptide chemistry, NMR data analysis, structure calculations. M.F.F.: antifungal and antibacterial assays. J.S.M. and K.J.R. project design and funding. C.D.P. K.J.R and M.F.F wrote the manuscript with input from J.S.M.

## Acknowledgements

This work was supported by Australian Research Council (ARC) grants DP120103369 and DP19190102058 to J.S.M. and K.J.R. C.D.P. was supported by a UQ postgraduate research award. K.J.R. and J.S.M were supported in part by ARC Future Fellowships FT130100890 and FT1201000013 respectively. This work utilized the Queensland NMR Network 900 MHz NMR spectrometer.

