## Supplementary Information for "Structural Characterization of the PawL-Derived Peptide Family, an Ancient Subfamily of Orbitides"

### Table of Contents:

**Supplementary Figure 1 |** MS and HPLC traces of all synthesized PLPs 2

**Supplementary Figure 2 |** Expansion of 1D $^{1}$H NMR spectra of PLPs. 3

**Supplementary Figure 3 |** Secondary 13Cα/13Cβ (black/grey) chemical shifts of PLPs. 4

**Supplementary Table 1 |** Temperature coefficients of HN protons of PLP peptides that were dissolved in 90:10 $H_2O/D_2O$ or 50:50 $H_2O/CD_3CN$. 5

**Supplementary Table 2 |** Defining angles of typical β-turns 6

**Supplementary Figure 4 |** Disc diffusion assay of PLPs against bacteria. 7

**Supplementary Figure 5 |** Disc diffusion assay of PLPs against *C. albicans*. 8

**Supplementary Table 3 |** Growth of *C. albicans* after 7 h in the presence of PLP-29 at various concentrations. 9

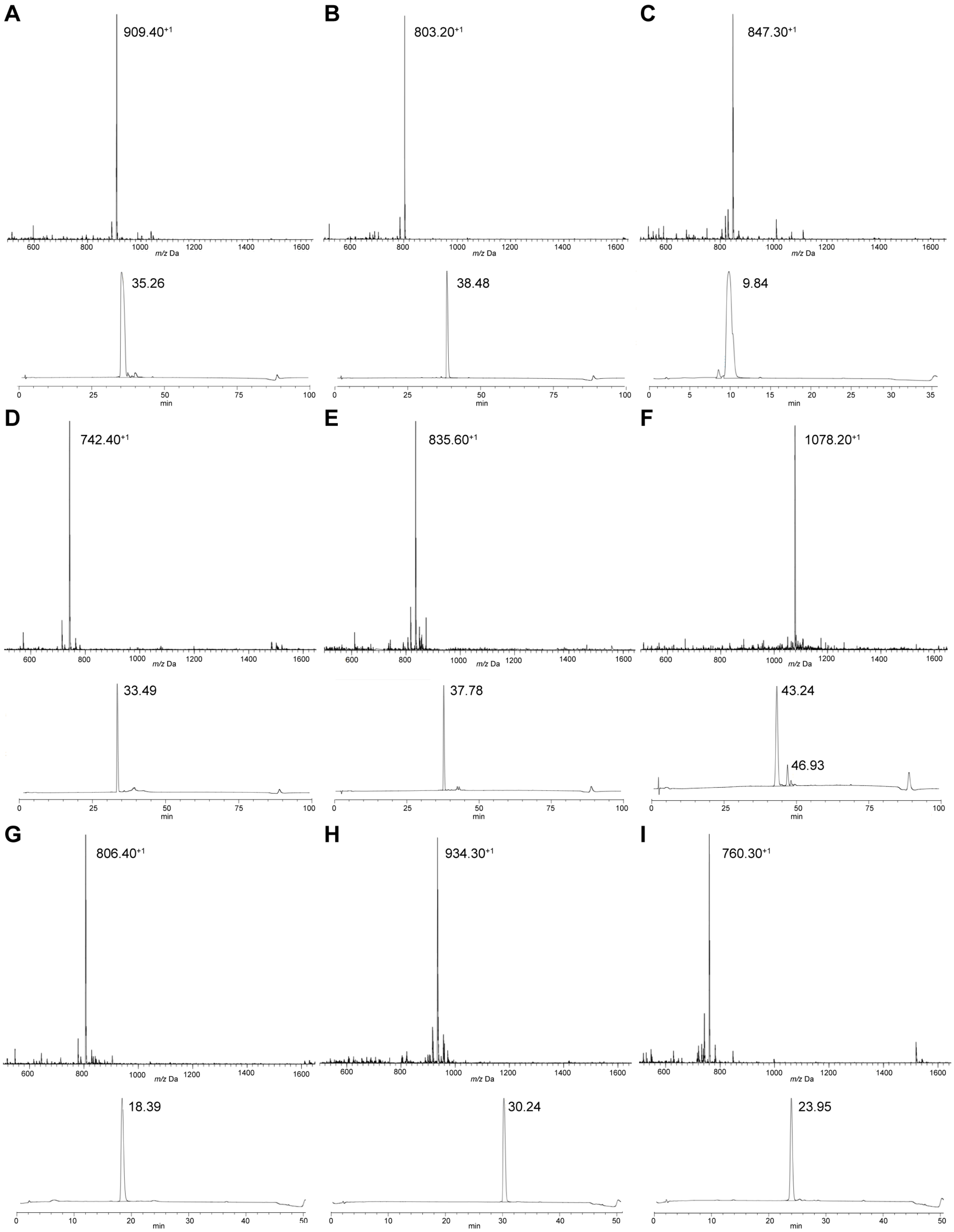


**Supplementary Figure 1 | MS and HPLC traces of all synthesized PLPs:** A-I: PLP-6, 13, -16, -22, -29, -31, -38, -42 and -46. Notably, the HPLC profile of PLP-31 contains two peaks, reflective of the presence of multiple conformations.

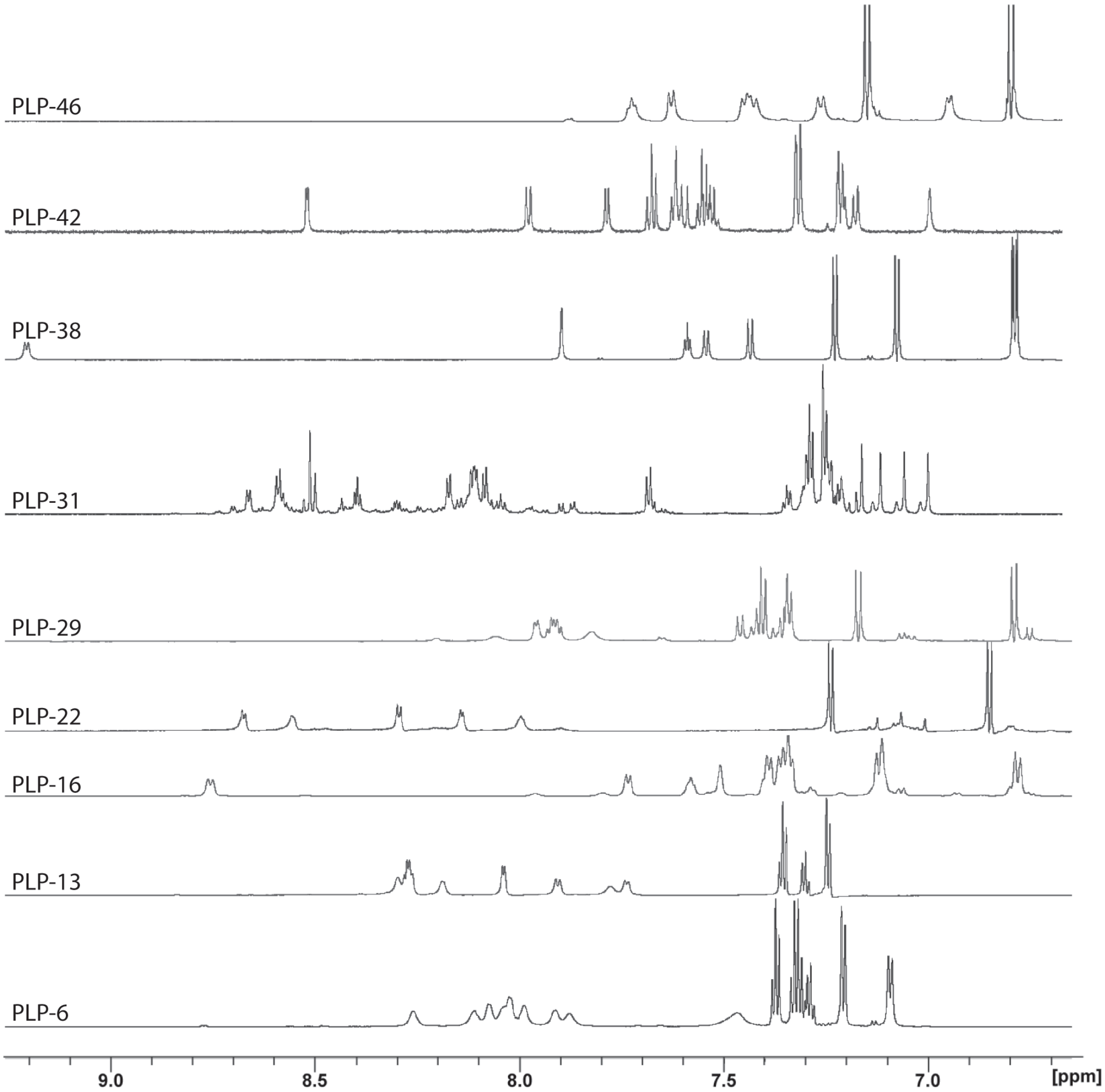


**Supplementary Figure 2 | Expansion of 1D 1H NMR spectra of PLPs.** All data were recorded at 900 MHz and 298 K.

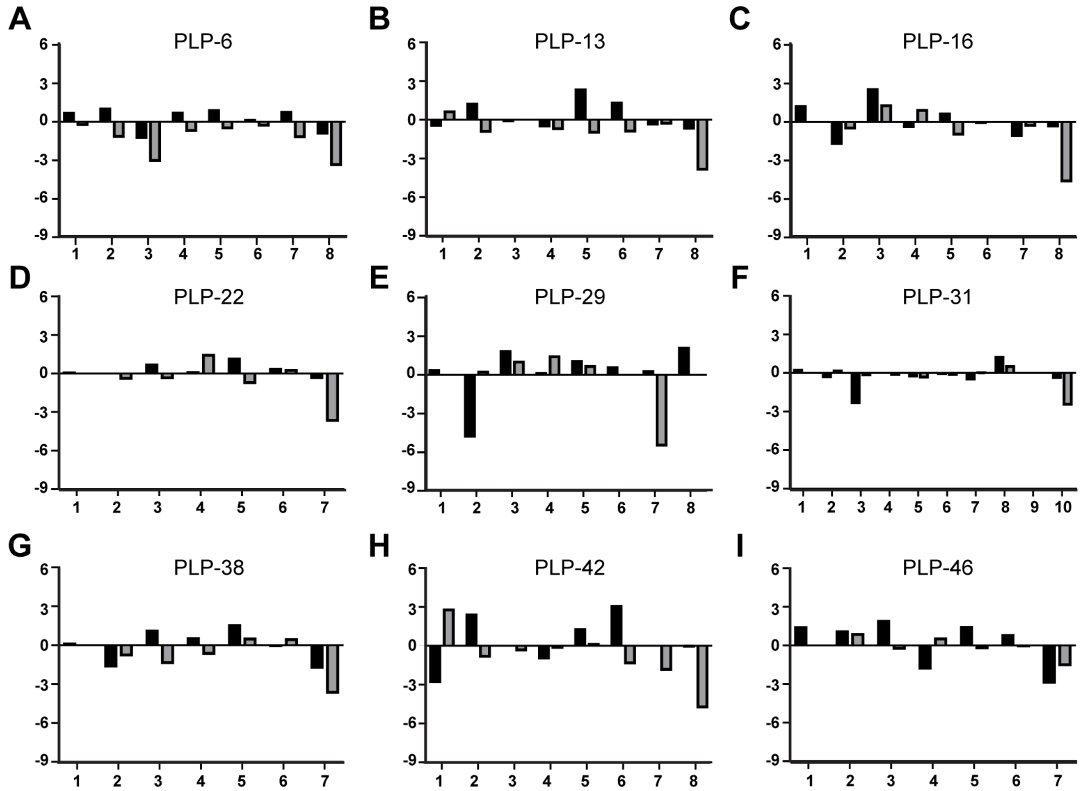


**Supplementary Figure 3 | Secondary 13Cα/13Cβ (black/grey) chemical shifts of PLPs.** A-I: PLP-6, -13, -16, -22, -29, -31, -38, -42 & -46. The y-axis for all graphs is the secondary chemical shift deviation (δΔ; ppm), X-axis is the residue number. Strong deviations are considered to be > 1 ppm.

| | Res 1 | Res 2 | Res 3 | Res 4 | Res 5 | Res 6 | Res 7 | Res 8 |
|---|---|---|---|---|---|---|---|---|
| PLP-13 | -2.8 | -8.0 | -7.5 | -5.6 | -6.8 | -11.8 | -2.3 | -3 |
| PLP-22 | -0.3 | -10.4 | Pro | Pro | -10.1 | -9.2 | -8.5 | |
| PLP-38 | -2.1 | -6.1 | -3.0 | Pro | N/A | Pro | -4.6 | |
| PLP-42 | -2.0 | -6.9 | -6.3 | -3.7 | Pro | -1.7 | -1.4 | -1.4 |

**Supplementary Table 1 | Temperature coefficients of HN protons of PLP peptides that were dissolved in 90:10 $H_2O/D_2O$ or 50:50 $H_2O/CD_3CN$.** Studies have shown that ~93% of amide protons with a temperature coefficient in water >-4.0 ppb/K are hydrogen bonded.[1] Yellow highlights identify amide protons that were found to be hydrogen bonded during structure calculation. The temperature data for PLP-13 and PLP-22, which were recorded in water are fully consistent with this observation and thus confirms the observed hydrogen bonding. Data recorded in 50:50 $H_2O/CD_3CN$ are however inconclusive for this type of analysis, as many amides showed a temperature dependence >-4.0 ppb/K but were not found to be hydrogen bonded during structure calculations. Proline residues, which lack a HN proton, are listed as Pro and N/A is used to denote a residue for which spin systems could not be resolved in the altered temperature NMR data. PLPs -22 and -38 only contain 7 residues.

**Supplementary Table 2 | Defining angles of typical β-turns**[2, 3]

| **β-turn** | $\phi_{i+1}$ | $\psi_{i+1}$ | $\phi_{i+2}$ | $\psi_{i+2}$ |
|---|---|---|---|---|
| I | -60 | -30 | -90 | 0 |
| I' | 60 | 30 | 90 | 0 |
| II | -60 | 120 | 80 | 0 |
| II' | 60 | -120 | -80 | 0 |
| III | -60 | -30 | -60 | -30 |
| III' | 60 | 30 | 60 | 30 |
| IV | Any turn differing by ≥ 2 angles from the list given | | | |
| V | -80 | 80 | 80 | -80 |
| V' | 80 | -80 | -80 | 80 |
| VIa[a] | -60 | 120 | -90 | 0 |
| VIb[a] | -120 | 120 | -60 | 0 |
| VII | Kink created by $\psi_2 \approx 180$ °, $\|\phi_3\| < 60$ °; or $\|\psi_2\|$, 60 °, $\phi \approx 180$ ° | | | |
| VIII | -60 | -30 | -120 | 120 |

[a] The peptide bond between the $i_{+1}$ and $i_{+2}$ residue is cis, and $i_{+2}$ is a Pro residue.

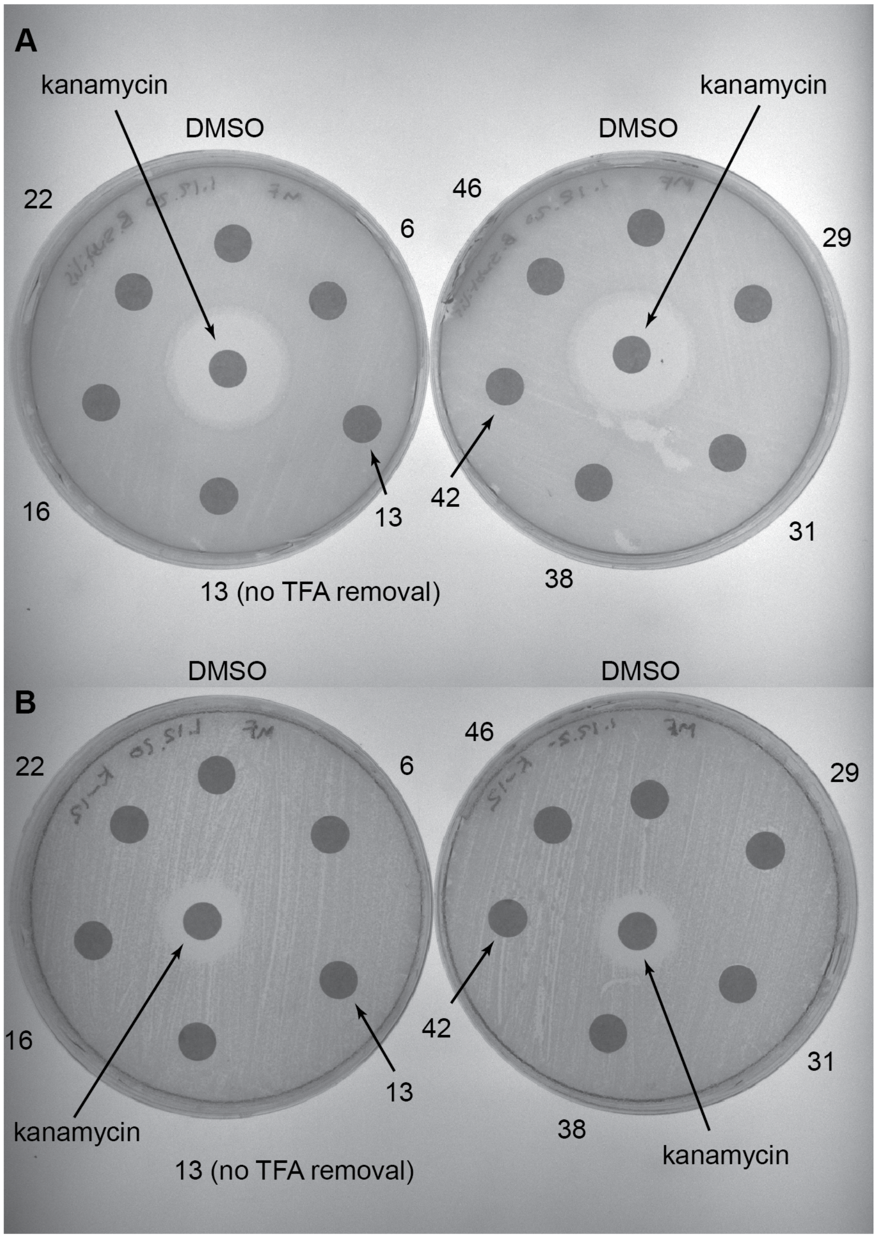


**Supplementary Figure 4 | Disc diffusion assay of PLPs against bacteria.** Discs containing 200 μg of each PLP were placed on LB agar plates inoculated with A: *B. subtilis* Marburg No. 165 or B: *E. coli* K-12. A disc containing DMSO was used as a negative control and a disc containing 50 μg of kanamycin was used as a positive control. PLPs are represented by their number. There was no indication of inhibition of bacterial growth, including for a TFA salt of PLP-13.

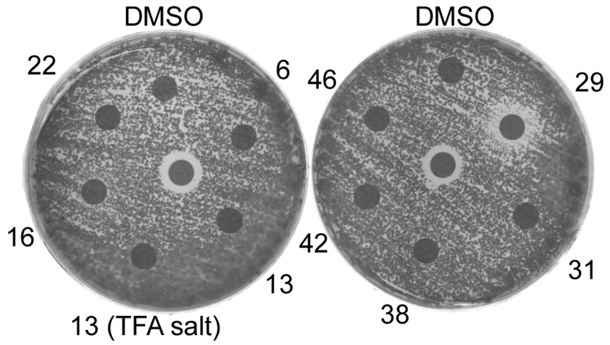


**Supplementary Figure 5 | Disc diffusion assay of PLPs against *C. albicans*.** Discs containing 200 μg of each PLP were placed on YPD agar plates inoculated with *C. albicans*. A disc containing DMSO was used as a negative control and a disc containing 50 μg of amphotericin B (center) was used as a positive control. Most PLPs demonstrated no inhibition of fungal growth, although PLP-29 showed minor inhibition of *C. albicans*.

| | 5 µM | 10 µM | 20 µM | 40 µM | 80 µM | 160 µM | 320 µM | 640 µM |
|---|---|---|---|---|---|---|---|---|
| No inoculum | 0.02 | 0.00 | 0.00 | 0.00 | 0.00 | 0.00 | -0.01 | -0.01 |
| DMSO Control | 0.85 | 0.92 | 0.85 | 0.92 | 0.86 | 0.94 | 0.78 | 0.73 |
| PLP-29 | 0.84 | 0.90 | 0.81 | 0.92 | 0.94 | 0.90 | 0.64 | 0.68 |
| PLP-29 | 0.84 | 0.88 | 0.83 | 0.89 | 0.84 | 0.87 | 0.71 | 0.69 |
| PLP-29 | 0.89 | 0.71 | 0.81 | 0.76 | 0.85 | 0.83 | 0.82 | 0.75 |
| Amphotericin B | 0.74 | 0.70 | 0.65 | 0.50 | 0.26 | 0.23 | 0.06 | 0.06 |
| Amphotericin B | 0.73 | 0.68 | 0.71 | 0.57 | 0.41 | 0.09 | 0.05 | 0.05 |
| Amphotericin B | 0.78 | 0.73 | 0.67 | 0.57 | 0.33 | 0.13 | 0.05 | 0.05 |
| Average (Mean) | | | | | | | | |
| Amphotericin B | 0.75 | 0.70 | 0.68 | 0.55 | 0.33 | 0.15 | 0.05 | 0.05 |
| PLP-29 | 0.86 | 0.83 | 0.82 | 0.86 | 0.87 | 0.87 | 0.72 | 0.71 |

**Supplementary Table 3 | Growth of *C. albicans* after 7 h in the presence of PLP-29 at various concentrations.** Three replicates of both PLP-29 and the amphotericin B positive control were used. Other controls were a "no inoculum" control containing growth medium and PLP-29 only and a DMSO control where DMSO was added to the fungal culture in the same concentrations as in the wells containing PLP-29.
